# Isolation of Pure Disease Specific Aging Trajectories in Spatial Transcriptomics via the Delta–Delta Method

**DOI:** 10.1101/2025.10.01.679749

**Authors:** Moonseok Choi, Sungsu Hwang, Kyu-Sung Kim, Jino Choi, Kyemyung Park, Do-Geun Kim

## Abstract

Disentangling normal aging from disease driven transcriptional change remains a major obstacle for spatial genomics. We introduce the Delta–Delta (ΔΔ) Method, a contrastive trajectory framework that resolves a four-dimensional progression (genes × cell types × brain regions × time) by subtracting the wild type (WT) aging trajectory from the transgenic (TG) trajectory to yield a pure disease trajectory (ΔΔlog₂FC). The method is platform agnostic, integrates with common spatial transcriptomics workflows, and outputs direction and speed of change summaries, enriched pathways, and region and cell type specific maps. In a demonstration using G2–3 α-synuclein TG mice and age matched WT controls at 6 and 10 months across hippocampus and midbrain, ΔΔ uncovered opposite regional dynamics in glutamatergic neurons and a convergent enrichment of RNA splicing pathways, corroborated by alternative splicing analyses. By explicitly modeling time while controlling for aging within each region and cell type, the ΔΔ Method isolates disease specific molecular programs that are obscured in conventional bulk or single cell analyses, and provides a generalizable framework for trajectory aware mechanistic target prioritization in neurodegeneration and other progressive conditions.

## Main

Neurodegenerative disorders such as Parkinson’s disease (PD) arise from complex, spatially heterogeneous molecular changes that unfold across brain regions and cell types over time ^1–3^. Conventional bulk RNA-seq lacks spatial and cellular resolution, obscuring region specific trajectories of decline and compensation ^4, 5^. Although single cell and single nucleus RNA-seq improve cell type specificity, they disrupt native tissue architecture and provide limited capacity to resolve temporal dynamics in situ ^6–8^. Spatial transcriptomics overcomes these constraints by enabling high throughput gene expression profiling in anatomically preserved sections, retaining both spatial coordinates and cellular context ^4, 9^.

Here we introduce “the Delta–Delta (ΔΔ) Method”, a contrastive trajectory framework that disentangles normal aging from disease driven transcriptional change while preserving spatial and cellular resolution. Coupled with high resolution spatial transcriptomics, ΔΔ resolves a four-dimensional progression (genes × cell types × brain regions × time) by subtracting the wild-type (WT) aging trajectory from the transgenic (TG) trajectory to yield a pure disease trajectory (ΔΔ log₂FC). The method is platform agnostic and integrates seamlessly with existing spatial transcriptomics workflows to generate direction and speed of change summaries, enriched pathway annotations, and region and cell type specific molecular maps.

To demonstrate its utility, we applied ΔΔ to G2–3 α-synuclein transgenic mice and age matched WT controls at 6 and 10 months, capturing both disease specific and normal aging processes in hippocampus and midbrain regions with differential vulnerability to neurodegeneration. ΔΔ analysis revealed opposite regional kinetics in glutamatergic neurons deceleration in midbrain and acceleration in hippocampus and highlighted RNA splicing pathways with representative factors (*Tardbp*, *Srsf3*, *Prpf8*, *U2af2*) corroborated by alternative splicing (AS) analyses ^10–13^.

By explicitly modeling time while controlling for aging within each region and cell type, the ΔΔ Method reframes spatial transcriptomics from static differential expression toward trajectory aware, aging controlled inference. This generalizable framework can be applied to diverse neurodegenerative and systemic disorders and extended to additional omics layers, offering a scalable approach to isolate disease specific molecular programs and support trajectory aware mechanistic target prioritization.

## Results

### Spatial transcriptomic annotation of brain regions and cell types in G2-3 mouse brain sections

We performed high resolution spatial transcriptomics on paraffin embedded brain sections from G2– 3 transgenic (TG) and age matched WT mice at 6 and 10 months using 10x Genomics Visium HD (2 µm bins), enabling near single cell analysis. Each hemisphere yielded over 160 million RNA detection events across 350,000–400,000 bins, detecting over 17,000 genes per sample (Fig. 1a).

**Fig. 1.**
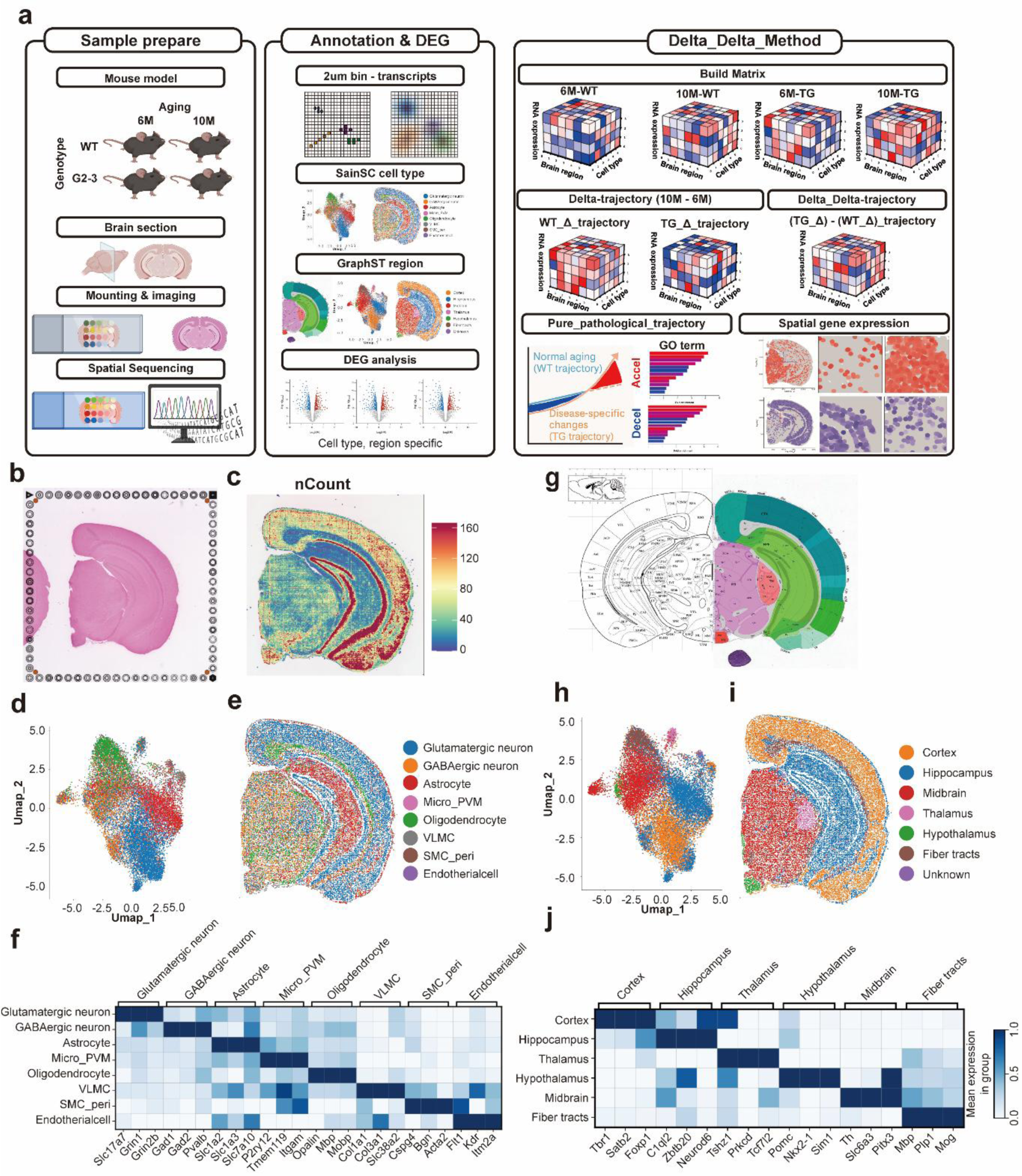
Spatial transcriptomics workflow and ΔΔ (Delta–Delta) analysis for pure trajectory inference. a, Experimental pipeline: sample preparation, 10x Genomics Visium HD capture, sequencing, and integration with the ΔΔ method to isolate disease specific transcriptomic trajectories. b, Representative H&E stained coronal section. c, Spatial distribution of mRNA abundance (nCount) across tissue as a quality control readout. d, UMAP of cell clusters annotated with SainSC using reference single cell datasets (Methods). e, Tissue map of annotated cell types, confirming expected spatial organization. f, Matrix plot of cell type marker genes showing distinct transcriptional signatures across major neuronal and glial populations. g, Brain atlas used as reference for GraphST based regional segmentation. h, UMAP colored by GraphST inferred anatomical regions. i, Tissue map of GraphST regional annotations showing concordance with known structures. j, Matrix plot of region specific marker genes validating segmentation across cortex, hippocampus, thalamus, hypothalamus, midbrain, and fiber tracts.

To preserve anatomical and cellular context, we implemented a dual layer annotation. Cell types were defined with SainSC guided by Allen Brain Atlas scRNA-seq references, consistently recovering major neuronal and glial classes (glutamatergic, GABAergic, dopaminergic, cholinergic neurons; astrocytes; oligodendrocytes; microglia; endothelial cells; VLMCs). Annotation accuracy was validated using canonical markers visualized by UMAP, dot plots, and cell-type marker matrices (Fig. 1b–f; Supplementary Fig. 1a, b, d–f)

Brain regions were delineated with GraphST ^14^, assigning spots to cortex, hippocampus, midbrain, thalamus, hypothalamus, and fiber tracts based on transcriptomic similarity to the Allen Mouse Brain Atlas; region calls were validated by region specific marker matrices and mapped consistently across WT and TG (Fig. 1g–j; Supplementary Fig. 1c–f).

Together, these results show that Visium HD near single cell spatial transcriptomics enables robust dual layer (cell type and regional) annotation, establishing a reproducible foundation for subsequent ΔΔ trajectory analyses.

### Region and cell type specific differential gene expression at each age in G2-3 mice

We profiled glutamatergic neurons from midbrain and hippocampus in TG and WT mice at 6 and 10 months. Volcano plots (cut off: |log₂FC| > 0.585; *p* < 0.05) revealed pronounced age × region differences (Fig. 2a,b). In the midbrain, glutamatergic neurons already showed robust alterations over 400 DEGs at with limited further expansion by 10 months, consistent with an early onset transcriptional shift. By contrast, the hippocampus exhibited few changes under 50 DEGs at 6 months but diverged strongly over 400 DEGs by 10 months, indicating delayed vulnerability relative to the midbrain.

**Fig. 2.**
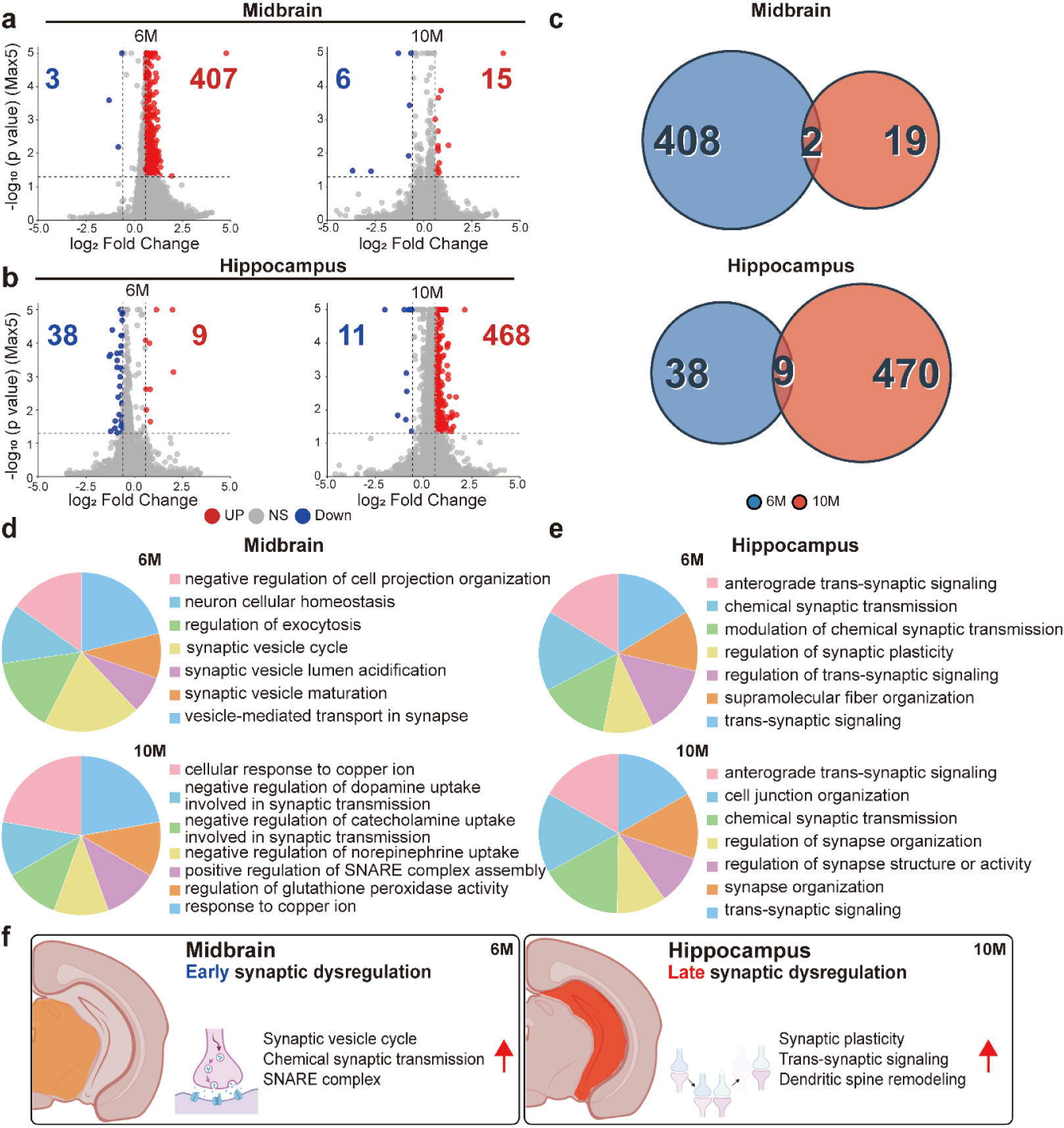
Region and age specific transcriptomic alterations in TG mice Glutamatergic neurons. a, b, Differential expression analysis (volcano plots) for glutamatergic neurons in midbrain (a) and hippocampus (b) at 6 and 10 months. Significance thresholds: |log₂FC| > 0.585 and *p* < 0.05 (red, up regulated; blue, down regulated; grey, not significant (NS)). c, Venn diagrams showing minimal overlap of significant transcripts between 6 and 10 month time points within each region. d, e, GOBP enrichment for region specific DEGs in midbrain (d) and hippocampus (e) at each age; top non redundant terms are shown (Methods). f, Schematic summary of the temporal regional cascade (early midbrain → late hippocampus); red arrows denote increased functional activity.

Venn analyses showed minimal overlap between 6 and 10 month DEGs within each region (Fig. 2c). GOBP enrichment further resolved these patterns: midbrain DEGs were enriched for synaptic vesicle cycle, chemical synaptic transmission, and trans synaptic signaling, whereas hippocampal DEGs at 10 months emphasized synaptic organization and plasticity related terms (Fig. 2d,e). To integrate across conditions, we summarized synapse associated processes among the top terms, revealing a convergent signature of synaptic organization, signaling, and vesicle dynamics (Fig. 2f).

Collectively, these results support a temporal regional cascade in glutamatergic neurons early synaptic dysregulation in midbrain (6 M) followed by late dysregulation in hippocampus (10 M) as encapsulated by the schematic panel (Fig. 2f). Analyses for GABAergic neurons, presented in (Supplementary Fig. 2), revealed a broadly similar early midbrain late hippocampus cascade, with early synaptic perturbations and delayed hippocampal changes paralleling the glutamatergic pattern. These observations motivated gene level interrogation of synaptic and activity dependent programs (Fig. 3).

**Fig. 3.**
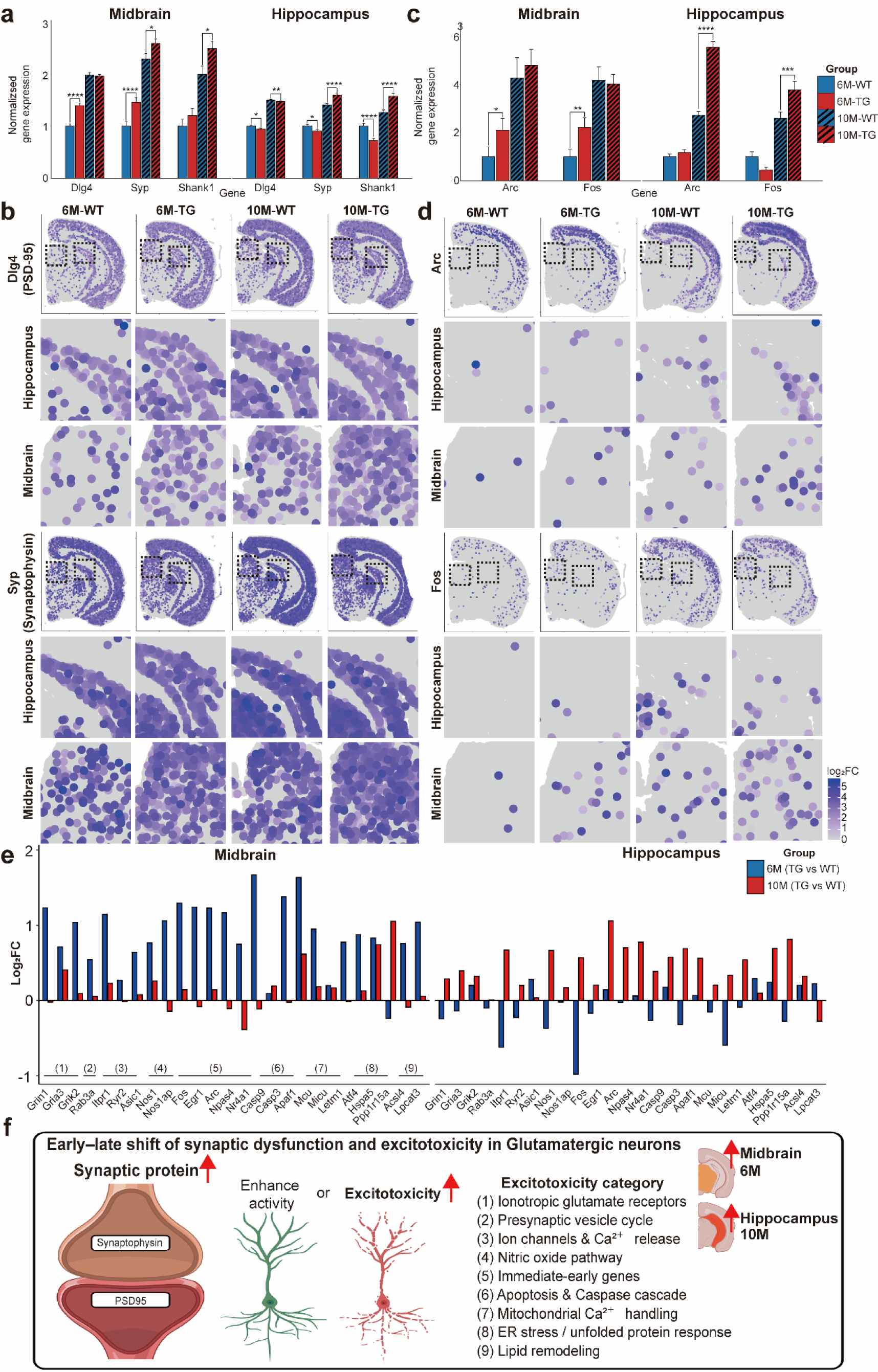
Spatially resolved analysis of synaptic and activity-dependent gene expression in Glutamatergic neurons. a, Bar graphs of synaptic marker genes (*Dlg4*/PSD-95, *Syp*/Synaptophysin, *Shank1*) in midbrain and hippocampus across four groups (6M-WT, 6M-TG, 10M-WT, 10M-TG), normalized to 6M-WT. Bars show mean ± S.E.; significance is indicated above bars (two-sided test with multiple-testing correction; details in Methods). b, Spatial transcriptomic maps (Visium HD) of *Dlg4* and *Syp* in tissue coordinates, displayed for each group and region. c, Bar graphs of immediate early genes (*Arc*, *Fos*) in the same format as (a), normalized to 6M-WT. d, Spatial maps of *Arc* and *Fos* expression in hippocampus and midbrain; expression hotspots localize to the superior colliculus motor-related (SCm) in midbrain and the dentate gyrus (DG) in hippocampus. e, Excitotoxicity related marker set plotted as log₂ fold-change (TG vs WT) at 6M (blue) and 10M (red), shown separately for midbrain and hippocampus. Representative genes span ionotropic glutamate receptors (e.g., *Grin1*, *Gria3*, *Grik2*), presynaptic vesicle cycle (*Rab3a*), Ca²⁺ channels/release (*Itpr1, Ryr2, Asic1*), nitric oxide signaling (*Nos1*, *Nos1ap*), immediate early genes (*Arc*, *Egr1*, *Fos*, *Nr4a1*, *Npas4*), apoptosis/caspase cascade (*Casp9, Casp3*, *Apaf1*), mitochondrial Ca²⁺ handling (*Mcu*, *Micu*, *Letm1*), ER stress/UPR (*Atf4*, *Hspa5*, *Ppp1r15a*), and lipid remodeling (*Acsl4, Lpcat3*). f, Schematic summary of the early late shift in Glutamatergic neurons: early synaptic dysregulation and heightened activity/excitotoxic drive in midbrain at 6M, followed by a late shift to hippocampus at 10M. Categories highlighted include (1) ionotropic glutamate receptors, (2) presynaptic vesicle cycle, (3) ion channels & Ca²⁺ release, (4) nitric-oxide pathway, (5) immediate early genes, (6) apoptosis & caspase cascade, (7) mitochondrial Ca²⁺ handling, (8) ER stress/UPR, and (9) lipid remodeling. Red arrows denote increased functional activity.

### Spatial transcriptomics reveal synaptic and neuronal activation programs in Glutamatergic neurons

At 2 µm (Visium HD) resolution, we quantified synaptic scaffolding genes (*Dlg4*/PSD-95, *Syp*/Synaptophysin, *Shank1*) and immediate early genes (IEG; *Arc*, *Fos*) ^15–19^ in Glutamatergic neurons of midbrain and hippocampus. Bar plot analyses (normalized to 6M-WT) showed significant increases in midbrain at 6 months, with no detectable change in hippocampus at 6 months; by 10 months, hippocampal glutamatergic neurons displayed robust upregulation (Fig. 3a,c). Spatial maps localized peak signals to SCm (midbrain) and DG (hippocampus), respectively (Fig. 3b,d).

To probe vulnerability pathways, we evaluated an excitotoxicity marker set spanning ionotropic glutamate receptors (*Grin1*, *Gria3*, *Grik2*) ^20–22^, Ca²⁺ handling/release (*Itpr1*, *Ryr2*, *Mcu*, *Micu2*, *Letm1*) ^23–26^, apoptotic mediators (*Casp3*, *Casp9*, *Apaf1*) ^27, 28^, lipid remodeling (*Acsl4*, *Lpcat3*) ^29, 30^, ER stress associated factors (*Atf4*, *Hspa5*, *Ppp1r15a*) ^31–33^, IEGs (*Fos*, *Egr1*, *Arc*) ^33, 34^, and nitric oxide signaling (*Nos1*, *Nos1ap*) ^35–37^. Glutamatergic midbrain showed a coordinated increase at 6 months, whereas the hippocampus exhibited minimal change at 6 months but marked elevation at 10 months (Fig. 3e). The schematic integrates these observations as an early midbrain, late hippocampus shift in synaptic/activation and excitotoxic drive (Fig. 3f). Corresponding GABAergic results, which recapitulate the early midbrain/late hippocampus motif with greater enrichment of inhibitory synapse and neurotransmitter metabolism terms, are provided in (Supplementary Fig. 3).

Despite the G2–3 α-synuclein TG model, the number of significant mRNAs at single ages remained modest, suggesting that aging exerts a dominant transcriptomic influence and can mask disease specific signals in cross sectional contrasts. These observations motivated a trajectory aware analysis to decouple aging from pathology; accordingly, we next model longitudinal change and apply the ΔΔ framework to isolate the pure disease trajectory.

### Aging trajectories reveal dominant transcriptomic changes

We defined the WT trajectory as aging associated change in WT (WT Δlog₂FC) and the TG trajectory as aging associated change in TG (TG Δlog₂FC). Unlike cross sectional TG–WT contrasts at a single age, trajectory analyses in glutamatergic neurons revealed widespread age driven reprogramming, with about 5,000–9,000 significant genes per trajectory depending on region (Fig. 4a, b), indicating that aging imposes a stronger transcriptional burden than α-synuclein overexpression alone. Comparing WT and TG trajectories within glutamatergic neurons showed substantial but region dependent overlap (Fig. 4c), the hippocampus displayed broad concordance in age responsive genes, whereas the midbrain overlapped less and exhibited genotype divergent patterns. Consistently, GOBP analysis highlighted shared housekeeping processes in the hippocampus (e.g., intracellular protein transport, membrane localization, mRNA metabolism, post translational modification), but divergent biological processes in the midbrain: WT enriched signaling/chemical synaptic transmission/trans synaptic signaling, whereas TG preferentially enriched (neuron/cell) projection morphogenesis (Fig. 4d,e). A presence–absence matrix summarizes recurrent top 7 GOBP term across regions (Fig. 4f).

**Fig. 4.**
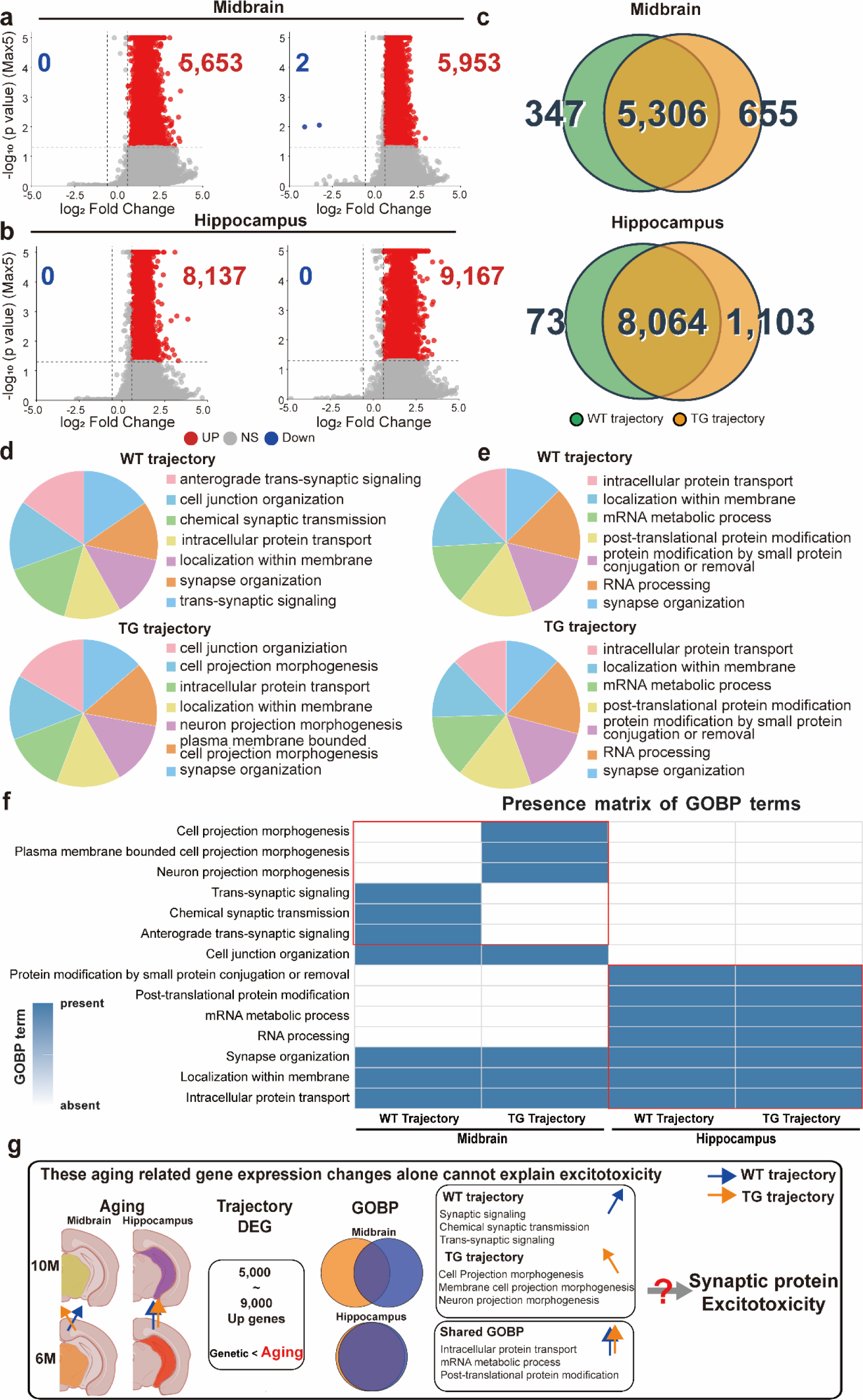
Within genotype aging trajectories in Glutamatergic neurons reveal region specific transcriptomic programs but are insufficient to explain excitotoxicity. a, b, Volcano plots of differentially expressed genes (DEGs) between 6 and 10 months computed within genotype (WT trajectory; TG trajectory) for glutamatergic neurons in midbrain (a) and hippocampus (b). Significance: |log₂FC| > 0.585; *p* < 0.05 (red, up regulated; blue, down regulated; grey, not significant (NS)). c, Venn diagrams showing overlaps of significant DEGs between WT and TG trajectories within each region. d, e, GOBP enrichment for trajectory DEGs in midbrain (d) and hippocampus (e), separated by genotype (Methods). f, Presence–absence matrix summarizing top recurrent synapse related GOBP terms across regions for WT and TG trajectories in glutamatergic neurons. g, Schematic of aging trajectory findings: aging within genotype yields widespread up regulation about 5,000–9,000 genes; hippocampus shows largely shared GOBPs between WT and TG, whereas midbrain diverges (WT: signaling/chemical synaptic transmission; TG: projection morphogenesis).

Despite the scale of age associated change, trajectory level enrichment does not by itself explain the observed excitotoxic signatures: term enrichment is list based and insensitive to direction and magnitude differences between WT and TG trajectories. These considerations motivated a disease specific correction. Accordingly, we applied the ΔΔ framework (ΔΔ = TG Δlog₂FC − WT Δlog₂FC) to isolate a pure disease trajectory that quantifies direction and speed of progression beyond aging (Fig. 4g), enabling mechanism focused analyses in subsequent sections. GABAergic trajectories (counts, overlaps, and GOBP enrichments) following the same pipeline are provided in (Supplementary Fig. 4) and recapitulate the early midbrain/late hippocampus motif with subtype specific nuances.

### Δ Δ Method isolates accelerated and decelerated transcriptomic trajectories

To disentangle aging related from disease specific dynamics, we quantified longitudinal change within each genotype: WT trajectory = WT Δlog₂FC (10 vs 6 months) and TG trajectory = TG Δlog₂FC. The pure disease trajectory was defined as ΔΔ = TG Δlog₂FC − WT Δlog₂FC, which tests whether disease accelerates (ΔΔ > 0) or decelerates (ΔΔ < 0) aging associated programs while preserving regional and cellular context. In glutamatergic neurons, matched heatmaps visualize WT, TG, and ΔΔ trajectories to capture both direction and speed of change across midbrain and hippocampus (Fig. 5a). Genes were classified as ΔΔ accelerated or ΔΔ decelerated based on the sign and significance/magnitude of ΔΔ relative to the WT aging baseline (rank/thresholding in Methods), yielding region specific gene sets (Fig. 5b).

**Fig. 5.**
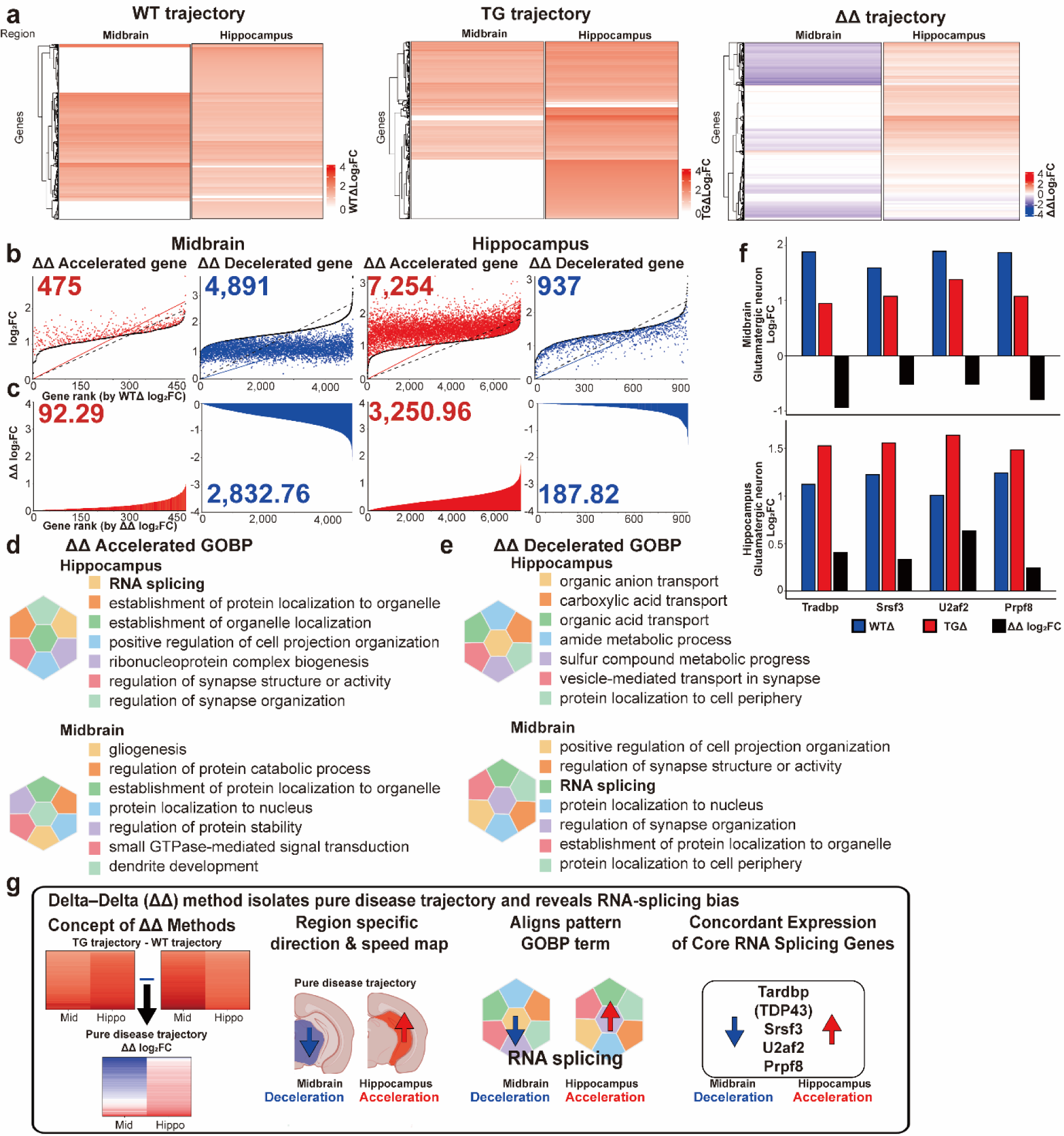
ΔΔ method isolates the pure disease trajectory and reveals regionally opposite RNA splicing dynamics in glutamatergic neurons. a, Trajectory heatmaps for WT (WT Δlog₂FC), TG (TG Δlog₂FC), and the pure disease trajectory (ΔΔ log₂FC = TG trajectory – WT trajectory) across midbrain and hippocampus in glutamatergic neurons. b, Rank ordered scatter plots identifying ΔΔ accelerated (red) and ΔΔ-decelerated (blue) genes in each region. c, Integrated effect of ΔΔ changes visualized as cumulative (area under curve:AUC) ΔΔ log₂FC for accelerated (red) and decelerated (blue) gene sets in midbrain and hippocampus. Acceleration/deceleration denotes faster/slower expression change in TG relative to the WT aging trajectory. d, e, GOBP enrichment using ΔΔ gene sets: (d) accelerated gene set; (e) decelerated gene set (Methods). f, Representative RNA splicing core genes (*Tardbp*, *Srsf3*, *U2af2*, *Prpf8*) exhibit regionally opposite ΔΔ trajectories decelerated in midbrain and accelerated in hippocampus consistent with pathway level enrichment. Bars show WT trajectory (blue), TG trajectory (red), and ΔΔ values (black). g, Schematic of the ΔΔ framework: subtracting the WT aging trajectory from TG isolates a pure disease trajectory, yielding a direction and speed map characterized by midbrain deceleration and hippocampal acceleration that aligns with RNA splicing GOBP and concordant expression of core splicing genes.

The midbrain showed a predominance of ΔΔ decelerated over 5,300 genes, indicating that disease slows age related transcriptional processes in excitatory neurons, whereas the hippocampus exhibited ΔΔ-acceleration over 8,000 genes, consistent with a late stage speed up. Importantly, gene count differences did not linearly predict expression kinetics: cumulative (area type) ΔΔ summaries (AUC) revealed shifts of about 20–28-fold in the midbrain and about 8–15-fold in the hippocampus (Fig. 5c), underscoring that ΔΔ quantifies speed, not merely the tally of significant genes. Corresponding GABAergic results as midbrain deceleration over 5,700 genes, hippocampal acceleration over 2,400 genes are provided in (Supplementary Fig. 5).

Functional partitioning, GOBP applied to ΔΔ gene sets separated accelerated versus decelerated biology (Fig. 5d,e) such as ΔΔ accelerated genes converged on RNA splicing, synaptic organization, and neuronal projection terms, whereas ΔΔ decelerated genes emphasized intracellular protein transport, metabolic processes, and vesicle mediated signaling a pattern robust across regions. At the gene level, core splicing regulators (*Tardbp*, *Srsf3*, *U2af2*, *Prpf8*) exhibited regionally opposite ΔΔ trajectories decelerated in midbrain and accelerated in hippocampus consistent with pathway level enrichment (Fig. 5f). The schematic integrates these findings into a ΔΔ based direction and speed map: disease slows excitatory transcriptional dynamics in the midbrain yet accelerates them in the hippocampus, aligning pattern level enrichment with core gene dynamics (Fig. 5g).

### Spatially resolved analysis of RNA splicing gene trajectories

We applied the ΔΔ framework to RNA splicing programs in glutamatergic neurons of the midbrain and hippocampus. Four core splicing factors (*Tardbp*, *Srsf3*, *U2af2*, *Prpf8*) showed significant ΔΔ effects upregulated in 10 month TG relative to the WT aging baseline whereas WT trajectories alone did not show comparable increases (Fig. 6a). Spatial maps localized these changes to SCm (midbrain) and DG (hippocampus), coinciding with synaptic/IEG hotspots (Fig. 6b). Quantification of alternative splicing (AS) across retained intron (RI), alternative 5′ splice site (A5SS), alternative 3′ splice site (A3SS), mutually exclusive exons (MXE), and skipped exon (SE) revealed a temporal regional shift: total AS events are elevated in midbrain at 6 months and increase in hippocampus by 10 months (Fig. 6c). The schematic integrates gene level, spatial, and AS category readouts with the ΔΔ direction and speed map, identifying RNA splicing as a disease relevant axis in the ΔΔ analysis (Fig. 6g). Corresponding GABAergic results (bar plots and spatial maps for the same four genes) are provided in (Supplementary Fig. 6), where effects are modest relative to glutamatergic neurons, underscoring cell type specificity.

**Fig. 6.**
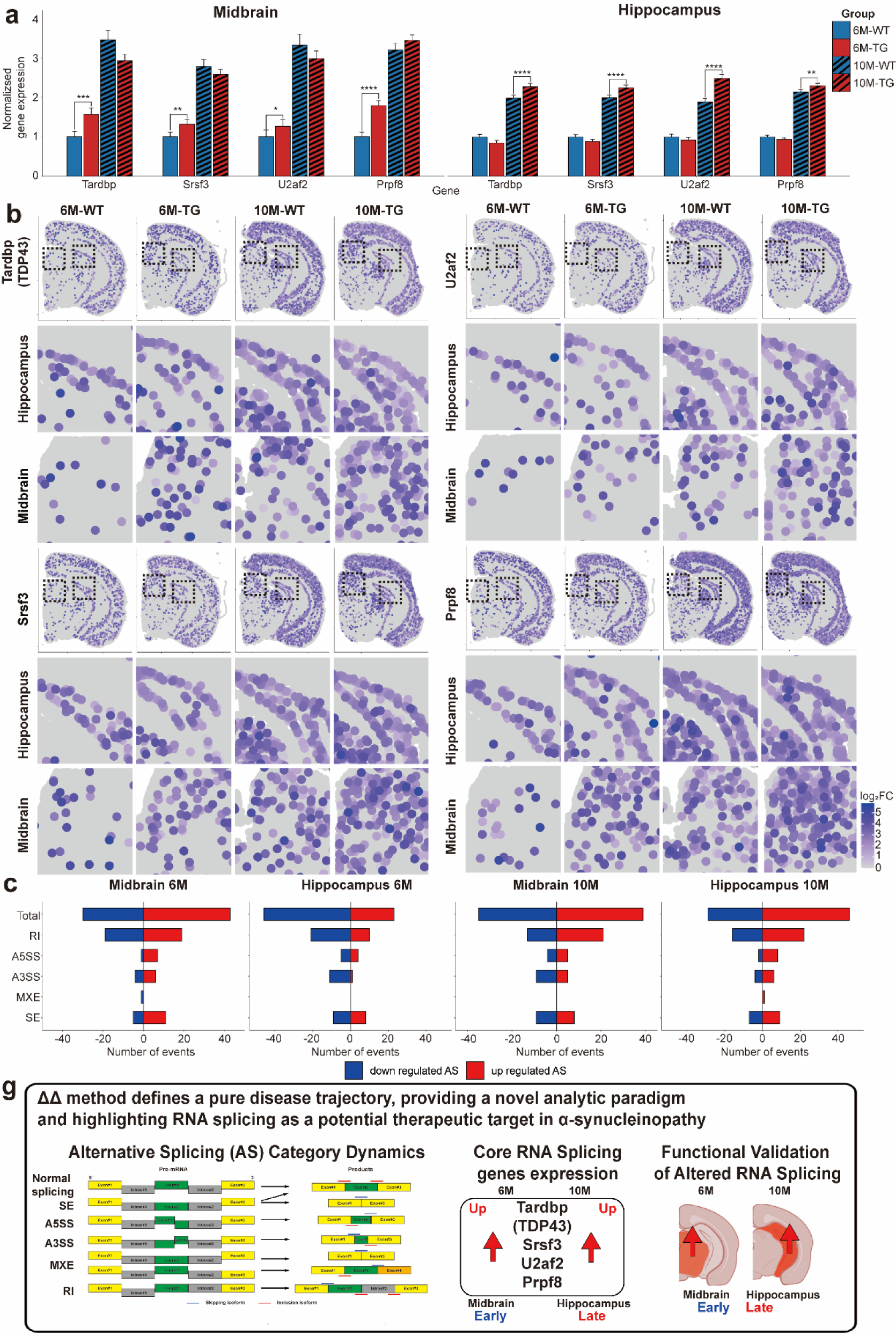
Spatially resolved analysis of RNA splicing gene expression and alternative splicing in Glutamatergic neurons. a, Bar plots showing normalized expression (relative to 6M-WT) of core RNA splicing genes (*Tardbp*, *Srsf3*, *U2af2*, *Prpf8*) in midbrain and hippocampus across four groups (6M-WT, 6M-TG, 10M-WT, 10M-TG). Bars show mean ± S.E. b, Spatial transcriptomic maps (10x Genomics Visium HD) depicting tissue distribution and expression intensity of the four RNA splicing genes in glutamatergic neurons across age and genotype; insets show high magnification fields for each region. c, Quantification of alternative splicing (AS) differences (TG vs WT) in midbrain and hippocampus at 6 and 10 months. Bars indicate the number of up regulated (red) and down regulated (blue) events for each AS category retained intron (RI), alternative 5′ splice site (A5SS), alternative 3′ splice site (A3SS), mutually exclusive exons (MXE), and skipped exon (SE) as well as the total number of AS events. g, Schematic summary. The ΔΔ framework (ΔΔ = TG trajectory – WT trajectory) defines a pure disease trajectory, revealing a region specific direction and speed map (early midbrain involvement → late hippocampus involvement). AS category dynamics together with concordant expression of core RNA splicing genes support altered RNA splicing as a disease relevant axis in α-synucleinopathy.

Together, these data indicate that ΔΔ methods defined, age modulated acceleration of RNA splicing processes contribute to the pure disease trajectory in vulnerable excitatory circuits. By explicitly controlling for aging, ΔΔ method resolves spatially specific splicing changes not evident in cross sectional DEGs and provides a general framework for trajectory aware target prioritization. Beyond PD, this framework is applicable to disorders with prominent temporal and spatial transcriptional dynamics (e.g., AD, HD, ALS, and cancer), and pure trajectories may inform translational biomarker development in peripheral biospecimens such as blood or CSF for early detection and monitoring of disease specific splicing dysfunction.

## Discussion

Neurodegenerative disorders such as Parkinson’s disease (PD) arise from spatiotemporally heterogeneous molecular perturbations that are not fully captured by conventional transcriptomic approaches ^38, 39^. Here we developed and applied the ΔΔ method in combination with high resolution spatial transcriptomics to resolve pure disease trajectories by subtracting the WT aging trajectory from the TG trajectory. This framework revealed cell type and region specific signatures most prominently synaptic dysfunction and excitotoxic stress that are not evident in cross sectional differential expression analyses.

A central observation is that aging exerts a stronger influence on global gene expression than α-synuclein overexpression in this model: trajectory based comparisons yielded about 2,600–9,100 significantly changing genes depending on region and cell type, far exceeding single age TG–WT contrasts. This aligns with clinical evidence that age is the dominant risk factor for PD ^40–42^, suggesting that genetic insults such as α-synuclein overexpression act in concert with age related vulnerabilities rather than as sole drivers. ΔΔ method further resolved differential susceptibilities across regions and neuronal subtypes: in midbrain, early changes were pronounced, whereas in hippocampus later changes predominated mirroring neuropathological staging in PD. These patterns were apparent only when aging trajectories were explicitly separated from transgene driven change. Across regions, synaptic scaffolding genes, IEGs, and excitotoxicity related genes increased along pathological trajectories, with SCm (midbrain) and DG (hippocampus) emerging as subregional hotspots, consistent with a shared excitotoxic component ^43, 44^. These results suggest a model whereby early synaptic hyperactivity and excitotoxic stress in select circuits contribute to vulnerability and subsequent degeneration.

Methodologically, ΔΔ method quantifies both the direction and speed of expression change while preserving anatomical and cellular context. Rather than static gene lists, the method generates a tensor (gene × cell type × region) populated by ΔΔ log₂FC, enabling classification of accelerated and decelerated genes and functional partitioning of these sets. This analysis highlighted common processes (e.g., synapse organization, intracellular transport) and, notably, RNA splicing as a disease associated, ΔΔ accelerated axis.

A key finding is the dysregulation of RNA splicing: core factors (*Tardbp*, *Srsf3*, *U2af2*, *Prpf8*) showed acceleration in TG trajectories especially in glutamatergic neurons supporting the broader view that aberrant splicing and RNA binding protein dysfunction cure across neurodegenerative diseases, including PD, AD, HD, and ALS ^45–47^. These changes were not observed in age matched WT trajectories, indicating disease specific rather than aging driven effects.

This framework generalizes conditions where aging and pathology intersect (e.g., AD, HD, ALS, and systemic diseases such as cancer) and can be extended to additional omics layers ^48–50^. The identification of splicing related alterations in tissue sections also motivates evaluation of peripheral biospecimens (blood, CSF) for translational biomarker development ^51–53^.

Several limitations warrant mention. First, Visium HD (2 µm bins) approaches but does not achieve single cell resolution; integration with single cell spatial platforms (e.g., MERFISH, Stereo-seq) will refine cell type specificity ^54, 55^. Second, our 50 bp read length constrains isoform level inference for certain alternative splicing events; longer reads (75–100 bp) or advanced platforms (e.g., Stereo-seq V2) should improve event detection ^56–58^. Third, GOBP enrichment was list based and did not weight expression magnitude, limiting sensitivity to quantitative differences. Fourth, splicing dysregulation was validated at the transcript level; protein level confirmation and functional assays remain important. Finally, two time points constrain modeling of non linear dynamics; denser sampling and larger cohorts will strengthen inference and reproducibility. Extensions to multi-omics and model based inference (including AI assisted regulator prediction) are natural next steps but require rigorous validation.

In summary, the ΔΔ method provides a method centric framework for spatial transcriptomics that resolves pure disease trajectories. Applied to a PD model, ΔΔ showed that aging outweighs α-synuclein overexpression in shaping transcriptional landscapes, uncovered region and cell type specific vulnerabilities, and identified excitotoxicity and RNA splicing as convergent disease relevant axes. Beyond PD, ΔΔ offers a scalable strategy for dissecting dynamic molecular processes across diseases, advancing mechanistic understanding and trajectory aware target prioritization.

## Methods

### Animals and Tissue Preparation

Male WT and G2–3 α-synuclein TG mice were maintained under identical housing on a 12h light/dark cycle with ad libitum access to food and water. Animals were euthanized at 6 and 10 months. After transcardial perfusion with PBS, brains were dissected, fixed in 4% paraformaldehyde for 24 h, paraffin embedded, sectioned coronally with a rotary microtome, and mounted on 10x Genomics Visium HD Gene Expression slides. Ethics. All procedures were approved by the institutional animal care and use committee (IACUC) and complied with relevant guidelines. (IACUC-23-00031/Korea Brain Research Institute: KBRI)

### Visium HD Spatial Gene Expression library

Formalin-fixed paraffin-embedded (FFPE) sections were processed per the manufacturer’s FFPE workflow (Visium HD; key documents CG000684/CG000685/CG000688). Briefly, sections were deparaffinized, H&E stained, imaged, and de-crosslinked. Probe hybridization and ligation were performed on-section; ligation products were transferred to the capture area via CytAssist, extended to append UMI, spatial barcode, and partial Read 1, and recovered for qPCR based cycle determination and sample indexing. Final libraries were purified (SPRIselect), quantified by qPCR (KAPA) and assessed on a Bioanalyzer (Agilent), then sequenced on Illumina NovaSeq using the read length recommended for Visium HD FFPE. The bin size for downstream analyses was 2 µm.

### Single cell Segmentation and Annotation using SainSC

To segment high-resolution data and assign cell types, we used SainSC (v0.5.0) ^59^. Lowly expressed genes were filtered (minimum count = 1). A Gaussian KDE (bandwidth = 4 µm) modeled spatial expression density; local maxima were detected (min distance = 4 pixels; min area = 60 pixels) to identify putative cell centers. Cell type annotation used a gene-signature matrix derived from the Allen Brain Atlas references (*Whole Cortex & Hippocampus SMART-seq 2019; 10x-SMART-seq taxonomy 2021*) ^60^ and cosine similarity between smoothed pixel profiles and signatures. A normalized assignment score enhanced separation between closely related subtypes. This yielded cell level annotation maps per section.

### Identification of Spatial Domains with GraphST

Spatial domains were identified with GraphST (v1.0.0) ^14^. Expression matrices were library-size normalized and log-transformed (SCANPY) ^61^ and the top 3,000 highly variable genes were selected. GraphST integrated expression and spatial coordinates to construct a graph neural representation; cells were partitioned into 20 clusters (Mclust; other parameters default) ^62^. Domains were manually annotated against the Allen Mouse Brain Atlas (cortex, hippocampus, midbrain, thalamus, hypothalamus, fiber tracts).

### Differential Expression Gene Analysis

To investigate transcriptional alterations related to genotype and age, DEG analysis was performed across four experimental groups based on two age points (6 and 10 months) and two conditions (WT and TG). The comparisons were conducted in two primary schemes within specific cell types and spatial domains. First, to identify genotype specific effects, DEG analysis was performed between WT and TG samples within the same age group (e.g., 6m-WT vs. 6m-TG). Second, to identify age related effects, DGE analysis was performed between the two age groups within the same genotype (e.g., 6m-WT vs. 10m-WT). For instance, glutamatergic neurons located within the ‘Cortex’ domain were compared across these four pairings.

We identified differentially expressed genes using Seurat’s FindMarkers() function (version 5.0.1), applying the default Wilcoxon rank-sum test (test.use = “wilcox”) to compare specified groups ^63, 64^. Genes were considered significantly differentially expressed if they met the criteria of an adjusted *p* value (*p* adj) < 0.05 and an absolute |log₂FC|> 0.585. The Benjamini-Hochberg procedure was used for multiple testing correction ^65, 66^. This approach allowed for the comprehensive identification of cell type and region specific transcriptional alterations driven by genotype and age.

### Trajectory Analysis

Mathematical Definition of Trajectories

g ∈ 𝒢: gene

c ɛ 𝒞: cell type

r ɛ ℛ: brain region

t ∈ {6, 10} (months): time

h ∈ {WT, TG}: genotype (WT; Wildtype, TG; Transgenic)

For each gene *g*, cell type *c*, brain region *r*, genotype h∈(WT, TG), and timepoint t∈(6M, 10M)t months, let

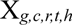

denote the normalized expression level of gene *g*.

**Step 1. Within-genotype aging trajectory (Δlog₂FC)**

The longitudinal change in gene expression within each genotype was calculated as the difference between 10-and 6-month expression levels:

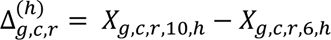

Stacking across all genes yields a trajectory vector for each region–cell type–genotype:

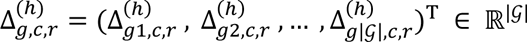

Here, Δ*_c,r_*^(WT)^ represents the WT trajectory, and Δ*_c,r_*^(TG)^ represents the TG trajectory.

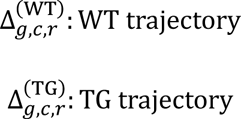

**Step 2. Pure trajectory (ΔΔlog₂FC)**

To isolate disease-specific changes, we defined the pure trajectory as the difference between TG and WT trajectories:

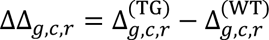

At the vector level:

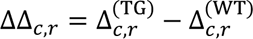

This forms a 3D matrix (*M*) indexed by (Z axes: expression - log_2_FC) × *c* (Y axes: GABAergic neuron, astrocyte and Glutamatergic) × *r* (X axes: Midbrain, Hippocampus and cortex).

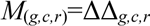

**Step 3. Intersection of genes**

To ensure comparability across genotypes, analyses were restricted to the intersection of genes detected in both WT and TG samples for the same region cell type:

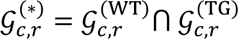

All trajectory quantities above use 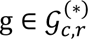

**Step 4. Acceleration vs deceleration and cumulative “speed”**

For each gene, disease specific trajectory speed was defined as:

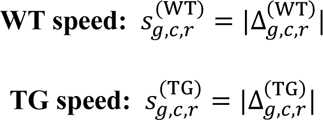

**Disease specific speed change** (your ΔΔ “speed”):

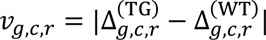

ΔΔ acceleration: *v_g,c,r_* > 0 (TG trajectory exceeds WT trajectory)

ΔΔ deceleration: *v_g,c,r_* < 0 (TG trajectory lags WT trajectory)

To summarize kinetics at the set level, we compute cumulative (area-type) summaries:

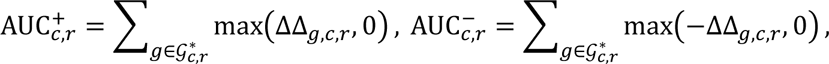

These capture the aggregate magnitude of ΔΔ acceleration and ΔΔ deceleration, respectively. The magnitude of the speed difference was quantified as:

This framework allows the systematic decomposition of age related transcriptional changes into WT trajectories, TG trajectories, and disease specific pure trajectories across brain regions and cell types.

### Functional Enrichment Analysis (GOBP)

clusterProfiler (v4.6.2) was used for GO Biological Process (BP) enrichment with org.Mm.eg.db ^67, 68^. For each brain region and cell type, DEG sets were defined with an adjusted *p* value (Benjamini– Hochberg) < 0.05 and an absolute log foldchange (|log_2_FC|) > 0.585. Up regulated and down regulated DEGs were analyzed separately. Gene symbols were mapped to Entrez Gene IDs using org.Mm.eg.db (Mus musculus), and enrichment was performed with enrichGO under the BP ontology ^69, 70^. The background gene universe was defined within each brain region and cell type, and terms with adjusted *p* < 0.05 (BH) were considered significant.

### Domain-specific alternative splicing (rMATS-turbo)

For a domain specific analysis, we first identified the cell barcodes corresponding to each anatomical region from an annotated object. Using these barcode lists, the primary alignment file was partitioned into domain specific BAM files with subset-bam (v1.1.0). Subsequently, we conducted a pseudo bulk differential alternative splicing analysis using rMATS turbo (v4.3.0) ^71–73^. This analysis compared corresponding spatial domains between age matched samples (e.g., 6 month WT vs. 6 month TG; 10 month WT vs. 10 month TG). The analysis was executed with the following parameters:-t single -- readLength 50--variable-read-length--allow-clipping. The output was then filtered to identify high-confidence differential splicing events across the compared conditions.

### Statistics

Unless stated otherwise, two-sided tests were used. Multiple testing was controlled by Benjamini– Hochberg FDR. For bar charts, bars represent mean ± SEM.

## Reporting summary

Further information on research design is available in the Nature Portfolio Reporting Summary linked to this article.

## Data availability

The datasets generated and analyzed in this study, including mouse model production, tissue sectioning, and 10x VisiumHD spatial transcriptomics, have not been deposited in a public repository as they have not been previously released. The data are available from the corresponding author upon reasonable request.

## Code availability

Custom R and Python scripts used for trajectory analysis, ΔΔlog₂FC computation, and functional enrichment are available at GitHub repository ^74^. Key packages include Seurat (v4.4.0) ^64^, clusterProfiler (v4.6.2) ^75^, GraphST ^14^, and SainSC ^59^. The exact parameters used for analysis are detailed in the Methods and Supplementary Information.

## Acknowledgements

We thank members of the Brain Research Institute for technical support and constructive discussions. We also acknowledge the 10x Genomics technical support team for assistance with Visium HD protocols.

## Author information

These authors contributed equally: Moonseok Choi, Sungsu Hwang

These authors jointly supervised this work: Do-Geun Kim, Kyemyung Park

Authors and Affiliations

**Dementia Research Group, Korea Brain Research Institute, Daegu, Republic of Korea**

Moonseok Choi, Kyu-Sung Kim, Jino Choi, Do-Geun Kim

**Graduate School of Health Science and Technology, Ulsan National Institute of Science and Technology (UNIST), Ulsan, Republic of Korea**

Sungsu Hwang, Kyemyung Park

**Department of Brain and Cognitive Sciences, Daegu Gyeongbuk Institute of Science and Technology (DGIST), Daegu, Republic of Korea**

Kyu-Sung Kim

**Department of Biomedical Engineering, UNIST, Ulsan 44919, Republic of Korea**

Kyemyung Park

## Contributions

M.C., K.P., and D-G.K. conceived and designed the study. M.C., S.H., K-S.K., and J.C. developed the methodology and analysis pipeline. M.C., and S.H. performed the investigations, including animal experiments, sequencing, and imaging. M.C., S.H., K-S.K., and J.C. carried out the formal data analysis, encompassing bioinformatics, statistical evaluation, and modeling. K.P. and D-G.K. provided resources and secured funding. M.C. wrote the original draft of the manuscript. All authors (M.C., S.H., K-S.K., J.C., K.P., and D-G.K.) contributed to reviewing and editing the manuscript. Visualization of figures and data plots was performed by M.C., S.H., K-S.K., and J.C. Supervision was provided by M.C., K.P., and D-G.K. Project administration was managed by K.P. and D-G.K.

## Corresponding authors

Correspondence to: Do-Geun Kim or Kyemyung Park.

## Supplementary information

Supplementary Figures (6).

