## Supplemental figures for "Isolation of Pure Disease Specific Aging Trajectories in Spatial Transcriptomics via the Delta–Delta Method"

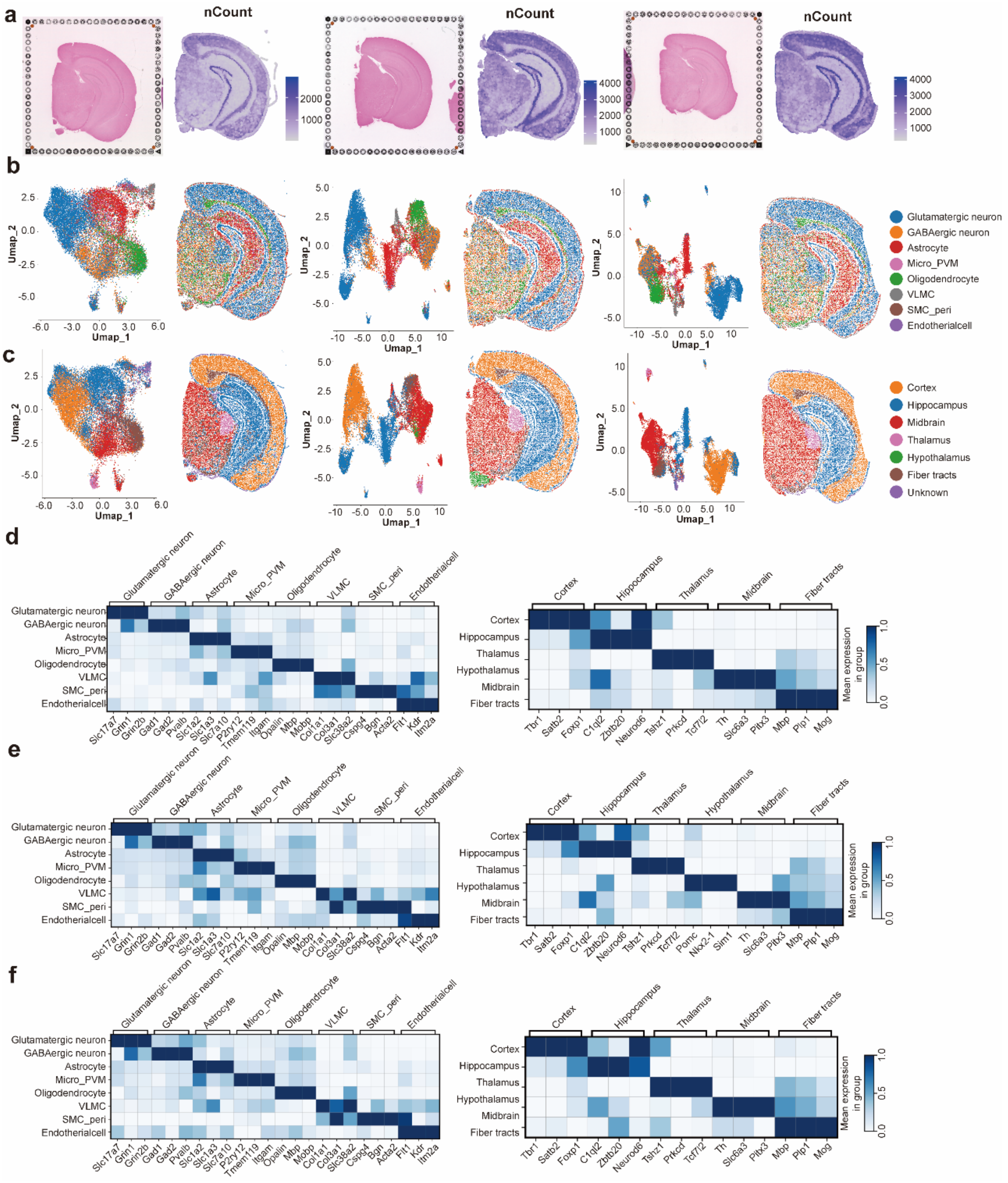


**Supplementary Fig. 1. Dual layer spatial annotation across experimental groups.**

a, Representative H&E-stained coronal sections for 6M-WT, 6M-TG, 10M-WT, and 10M-TG mouse brains. b, SainSC-based cell type annotation: UMAP projections and spatial maps showing classification into glutamatergic neurons, GABAergic neurons, astrocytes, oligodendrocytes, dopaminergic neurons, cholinergic neurons, microglia, endothelial cells, and VLMCs across all groups. c, GraphST-based region annotation: UMAP projections and spatial mapping of six anatomical domains (cortex, hippocampus, midbrain, thalamus, hypothalamus, fiber tracts). d–f, Matrix plots summarizing the top marker genes for each annotated cell type (left) and brain region (right), shown separately for (d) 6M-WT, (e) 10M-WT, and (f) 10M-TG groups.

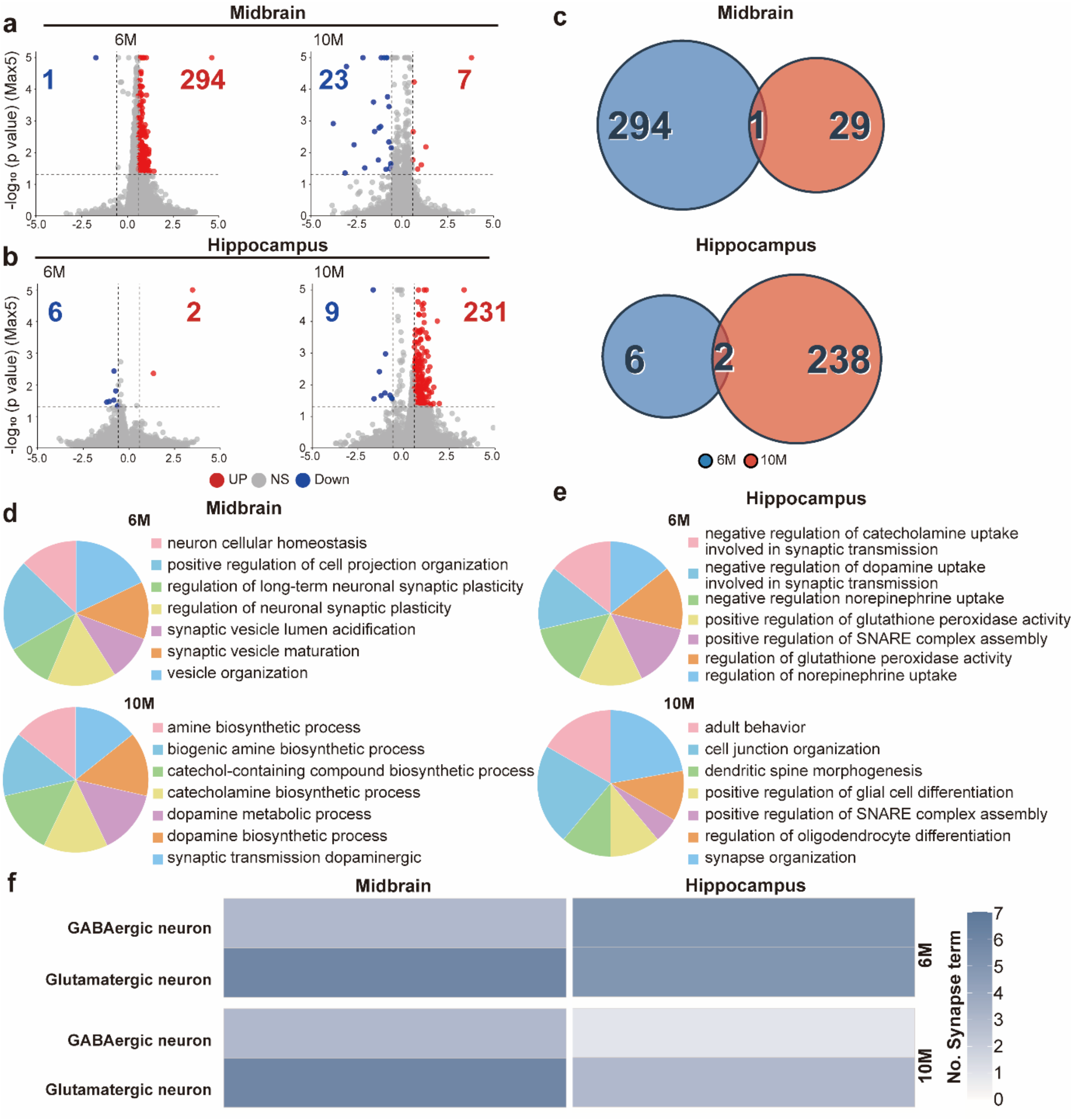


**Supplementary Fig. 2. Region and age specific transcriptomic alterations in TG mice GABAergic neurons.** (a, b), Differential expression analysis (volcano plots) of GABAergic neurons in the midbrain (a) and the hippocampus (b) at 6 and 10 months, highlighting significantly up and down regulated genes (cut off: $|\log_2 FC| > 0.585$, $p < 0.05$). c, Venn diagrams showing the overlap of significantly altered transcripts between 6 and 10 month time points within each region. (d, e), GOBP enrichment for midbrain (d) and hippocampus (e) region specific DEGs in GABAergic neurons at 6 and 10 months. f, Presence matrix summarizing the number of synapse related GOBP terms across regions and ages for glutamatergic neuron and GABAergic neurons.

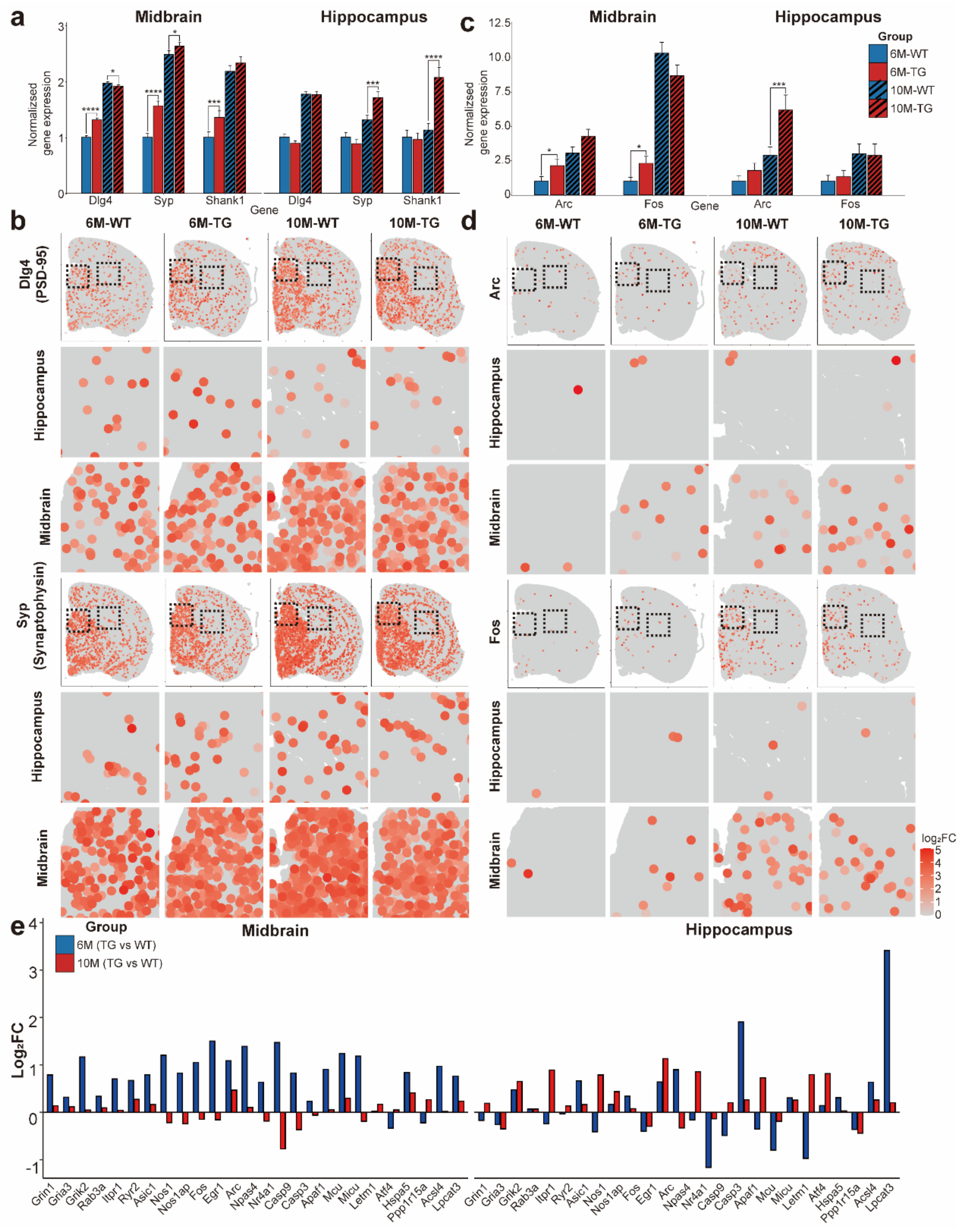


**Supplementary Fig. 3. Spatially resolved analysis of synaptic and activity dependent gene expression in GABAergic neurons.** a, Bar graphs of synaptic marker genes (*Dlg4*/PSD-95, *Syp*/Synaptophysin, *Shank1*) in midbrain and hippocampus across four groups (6M-WT, 6M-TG, 10M-WT, 10M-TG), normalized to 6M-WT. Bars show mean ± s.e.m.; significance is indicated above bars (two sided tests with multiple-testing correction; details in Methods). b, Spatial transcriptomic maps (Visium HD) of *Dlg4* and *Syp* plotted in tissue coordinates for each group and region; insets show high magnification fields. c, Bar graphs of immediate early genes (*Arc*, *Fos*) in the same format as (a), normalized to 6M-WT. d, Spatial maps of *Arc* and *Fos* expression in hippocampus and midbrain; expression hotspots localize to the superior colliculus motor related (SCm) in midbrain and the dentate gyrus (DG) in hippocampus. e, Excitotoxicity related marker set plotted as $\log_2$FC (TG vs WT) at 6M (blue) and 10M (red), shown separately for midbrain and hippocampus. Representative genes include ionotropic glutamate receptors (e.g., *Grin1*, *Gria3*, *Grik2*), presynaptic vesicle cycle (*Rab3a*), $Ca^{2+}$ channels/release (*Itpr1*, *Ryr2*, *Asic1*), nitric oxide signaling (*Nos1*, *Nos1ap*), IEGs (*Arc*, *Egr1*, *Fos*, *Nr4a1*, *Npas4*), apoptosis/caspase cascade (*Casp9*, *Casp3*, *Apaf1*), mitochondrial $Ca^{2+}$ handling (*Mcu*, *Micu*, *Letm1*), ER stress/UPR (*Atf4*, *Hspa5*, *Ppp1r15a*), and lipid remodeling (*Acsl4*, *Lpcat3*).

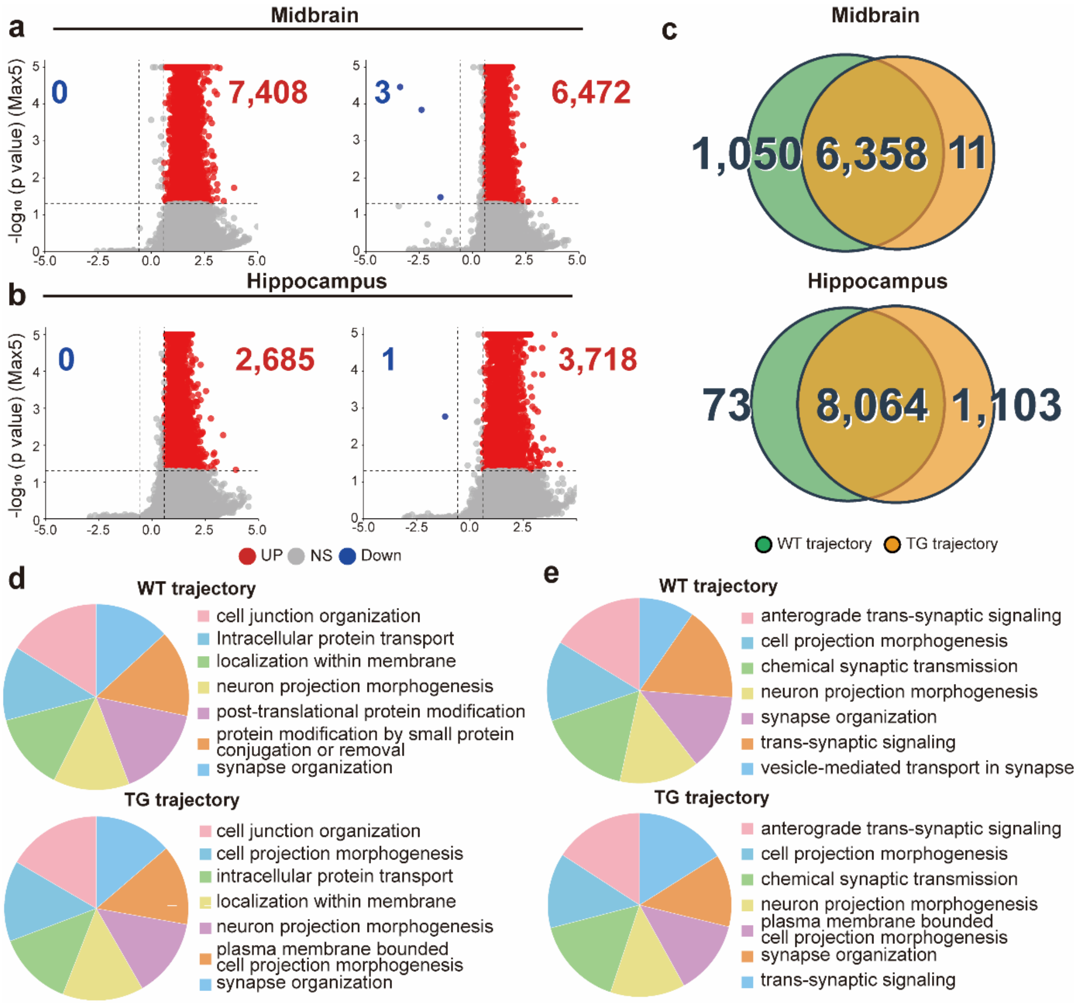


**Supplementary Fig. 4. Within genotype aging trajectories in GABAergic neurons reveal region specific transcriptomic programs.** (a, b), Volcano plots of differentially expressed genes (DEGs) between 6 and 10 months computed within genotype (WT trajectory; TG trajectory) for GABAergic neurons in the midbrain (a) and hippocampus (b). Points denote individual genes (red, up regulated; blue, down regulated; grey, not significant (NS)); numbers indicate significant genes. (cut off: $|log_2FC| > 0.585$, $p < 0.05$). c, Venn diagrams showing the overlap of significant DEGs between WT and TG trajectories within each region. (d, e), Gene Ontology Biological Process (GOBP) enrichment for trajectory DEGs in the midbrain (d) and hippocampus (e).

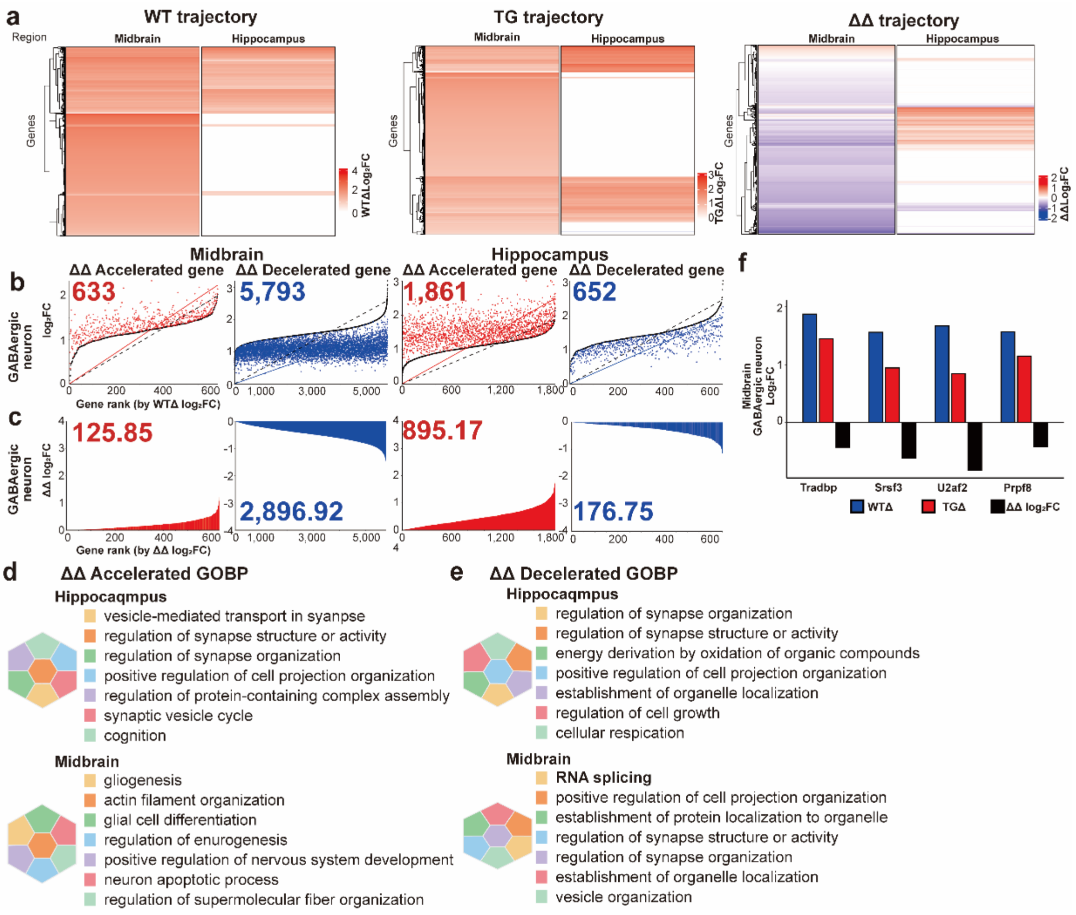


**Supplementary Fig. 5. ΔΔ method isolates the pure disease trajectory and reveals regionally opposite RNA splicing dynamics in GABAergic neurons.** a, Trajectory heatmaps for WT (WT $\Delta\log_2$FC), TG (TG $\Delta\log_2$FC), and the pure disease trajectory (ΔΔ $\log_2$FC = TG – WT) across midbrain and hippocampus in GABAergic neurons. b, Rank ordered scatter plots identifying ΔΔ accelerated (red) and ΔΔ decelerated (blue) genes in each region. c, Cumulative ΔΔ $\log_2$FC (area type summaries) quantifying the integrated effect of accelerated (red) and decelerated (blue) gene sets in midbrain and hippocampus. Acceleration/deceleration denotes faster or slower expression change in TG relative to the WT aging trajectory. (d, e), Gene Ontology Biological Process (GOBP) enrichment of ΔΔ gene sets. ΔΔ accelerated genes highlight processes such as vesicle mediated transport in synapse, regulation of synapse structure or activity, regulation of synapse organization, and synaptic vesicle cycle in the hippocampus, and gliosis, actin filament organization, and glial cell differentiation in the midbrain. ΔΔ decelerated genes are enriched for regulation of synapse organization and structure, organelle localization and cell growth in the hippocampus, and RNA splicing and related protein localization processes in the midbrain. f, Representative RNA-splicing core genes (*Tardbp*, *Srsf3*, *U2af2*, *Prpf8*) exhibit ΔΔ deceleration in the midbrain and comparatively stable or modest changes in the hippocampus, consistent with the pathway level GOBP enrichment. Bars show WT trajectory (blue), TG trajectory (red), and ΔΔ values (black).

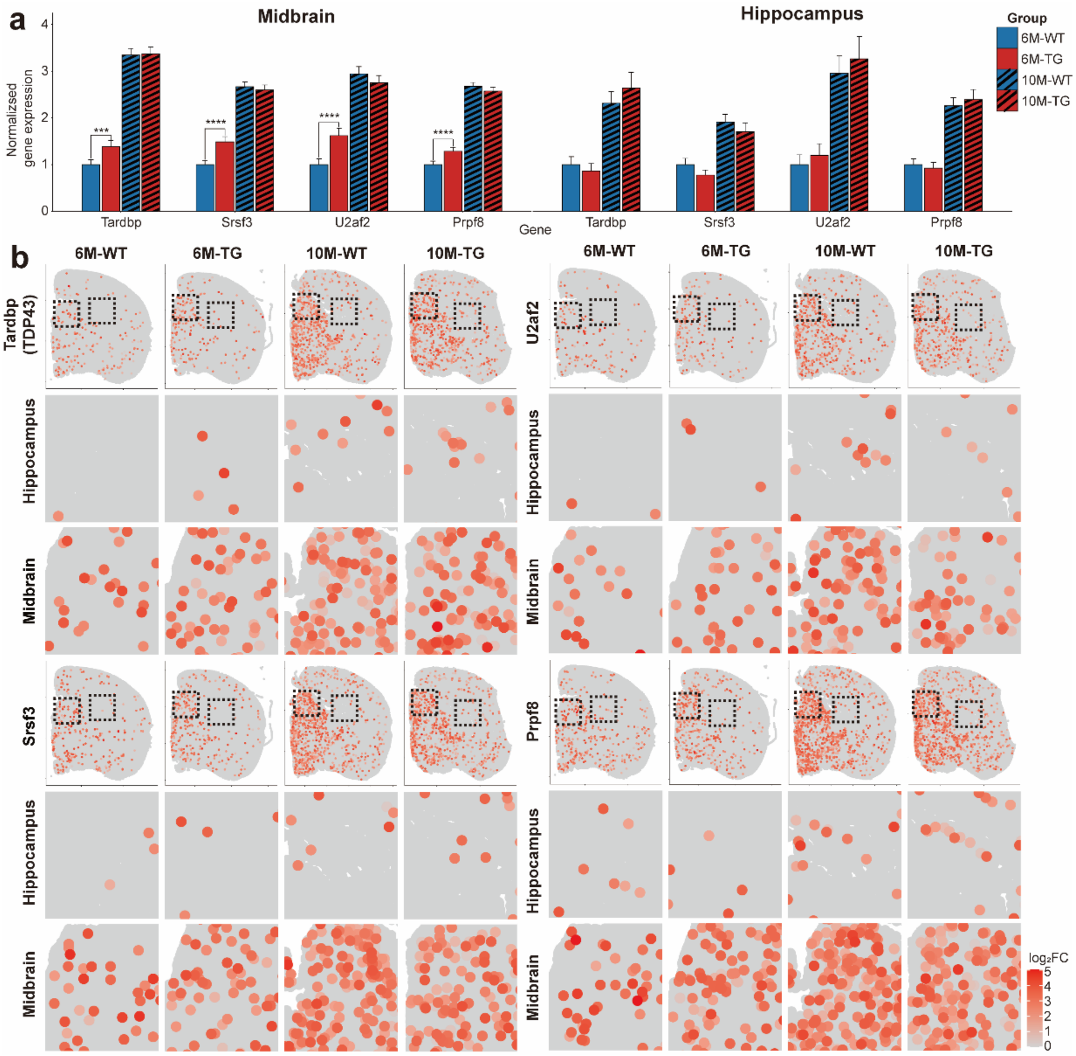


**Supplementary Fig. 6. Spatially resolved analysis of RNA splicing gene expression in GABAergic neurons.** a, Bar plots showing normalized expression (relative to 6M-WT) of core RNA splicing genes (*Tardbp*, *Srsf3*, *U2af2*, *Prpf8*) in midbrain and hippocampus across four groups (6M-WT, 6M-TG, 10M-WT, 10M-TG). Bars represent mean ± s.e.m. b, Spatial transcriptomic maps (Visium HD) depicting tissue distribution and expression intensity of these four RNA splicing genes in GABAergic neurons across age and genotype; insets show high-magnification views for each region.
